# Comparative Genomic Analysis of *Bacillus thuringiensis* Reveals Molecular Adaptations to Copper Tolerance

**DOI:** 10.1101/360354

**Authors:** Low Yi Yik, Grace Joy Wei Lie Chin, Collin Glen Joseph, Kenneth Francis Rodrigues

## Abstract

*Bacillus thuringiensis* is a type of Gram positive and rod shaped bacterium that is found in a wide range of habitats. Despite the intensive studies conducted on this bacterium, most of the information available are related to its pathogenic characteristics, with only a limited number of publications mentioning its ability to survive in extreme environments. Recently, a *B. thuringiensis* MCMY1 strain was successfully isolated from a copper contaminated site in Mamut Copper Mine, Sabah. This study aimed to conduct a comparative genomic analysis by using the genome sequence of MCMY1 strain published in GenBank (PRJNA374601) as a target genome for comparison with other available *B. thuringiensis* genomes at the GenBank. Whole genome alignment, Fragment all-against-all comparison analysis, phylogenetic reconstruction and specific copper genes comparison were applied to all forty-five *B. thuringiensis* genomes to reveal the molecular adaptation to copper tolerance. The comparative results indicated that *B. thuringiensis* MCMY1 strain is closely related to strain Bt407 and strain IS5056. This strain harbors almost all available copper genes annotated from the forty-five *B. thuringiensis* genomes, except for the gene for Magnesium and cobalt efflux protein (CorC) which plays an indirect role in reducing the oxidative stress that caused by copper and other metal ions. Furthermore, the findings also showed that the Copper resistance gene family, CopABCDZ and its repressor (CsoR) are conserved in almost all sequenced genomes but the presence of the genes for Cytoplasmic copper homeostasis protein (CutC) and CorC across the sample genomes are highly inconsonant. The variation of these genes across the *B. thuringiensis* genomes suggests that each strain may have adapted to their specific ecological niche. However, further investigations will be need to support this preliminary hypothesis.

## 1. Introduction

Copper is an important trace element required by living organisms in a very small amount to sustain the functioning of enzymes. However, excessive amount of copper can be harmful to life, this is because of its semi-soluble property in water. Copper can perform one electron redox chemistry and produces free Cu(I) radicals that target the enzymes associated with the intermediary metabolism [1]. In nature, copper occurred in rock, soil or sediment systems, due to the disturbance of natural event or human activities such as mining operations, overexposure of copper minerals occurs and this can lead to environmental pollutions and the chain reaction can affect the ecosystem. Some microorganisms such as bacteria are known to have high endurance towards copper pollution and able to thrive in extreme niches which are contaminated by copper [2]. *B. thuringiensis* is a rod shaped soil bacteria that have also been studied intensively as a candidate of biological pesticide. In early of the 20^th^ century, *B. thuringiensis* was discovered as a pathogen to silk worm larvae and it was named after a German Town, Thuringia later [3]. After several decades of investigations, the pathogenicity of *B. thuringiensis* was found to be associated with the Crystal protein produced by the bacteria itself. Over the years, several publications had made to prove that *B. thuringiensis* were found to be surviving at environments that are associated with copper pollution, such as acid mine drainage [4], heavy metal polluted wastewater [5] and industrial effluent contaminated soil [6]. However, some publications also addressed this bacteria as a common mesophile that has minimal tolerance to copper [7]. Nevertheless, none of these studies have provided any comparisons in the genetic and molecular aspects. Recently, a *Bacillus thuringiensis* strain MCMY1 was isolated from the copper contaminated soil collected from Mamut Copper Mine, Sabah, Malaysia [8]. The genome of *B. thuringiensis* MCMY1 was sequenced by using Single Molecular Real Time sequencing (SMRT) approach of Pacific Bioscience and assembled by using SMRT Portal. *B. thuringiensis* strain MCMY1 had shown its ability to withstand a Minimum Inhibitory Concentration (MIC) of 1.7 mM of simulated copper stress in previous experiment [8]. This study is driven by the interest to investigate the evolutionary adaptation of *B. thuringiensis* to copper tolerance by using comparative genomic approach. *B. thuringiensis* MCMY1 strain was used as the target strain for comparison with other available completed genomes of *B. thuringiensis* in NCBI GenBank database.

## 2. Materials and Methods

The genome of *B. thuringiensis* strain MCMY1 (PRJNA374601) was used as the target genome for comparison analysis. Other *B. thuringiensis* genomes were selected among those submitted as of May, 2018 in NCBI GenBank genome database and together with a control genome of *Escherichia coli* strain O157:H7 Sakai. All *B. thuringiensis* genomes were selected by referring to the genome and assembly report of *B. thuringiensis* under the category of completed genome in the GenBank. The list of all *B. thuringiensis* genomes and their responding strains have been described in Table 1. The nucleotide sequences of all strains were retrieved in FASTA format from the GenBank genome database. The quality of the assemblies for all genomes were assessed by using QUAST version 4.6.3 [9]. Whole genome alignment and Fragment all-against-all comparison analysis was done by using Gegenees software version 2.2.1 with BLAST+ version 2.7.1 installed in the pipeline [10]. A heat plot for the genomes comparison was generated from Gegenees software and exported in Nexus file format to be used for further phylogenomic analysis. SplitsTree software version 4 [11] was used for computing of phylogenetic tree. All *B. thuringiensis* genomes selected were uploaded to Rapid Annotations using Subsystems Technology (RAST) version 2.0 for annotations [12]. The summary of all annotations were viewed in the SEED viewer of RAST [13]. All genes annotated in the subsystems of copper homeostasis, copper tolerance, copper uptake, copper transport system and blue copper proteins were retrieved from the database. BLAST Ring Image Generator (BRIG) was used to compare the presence of these annotated copper related genes with a customized multi-FASTA file that contained the annotations information as the reference genome [14].

**Table 1.**
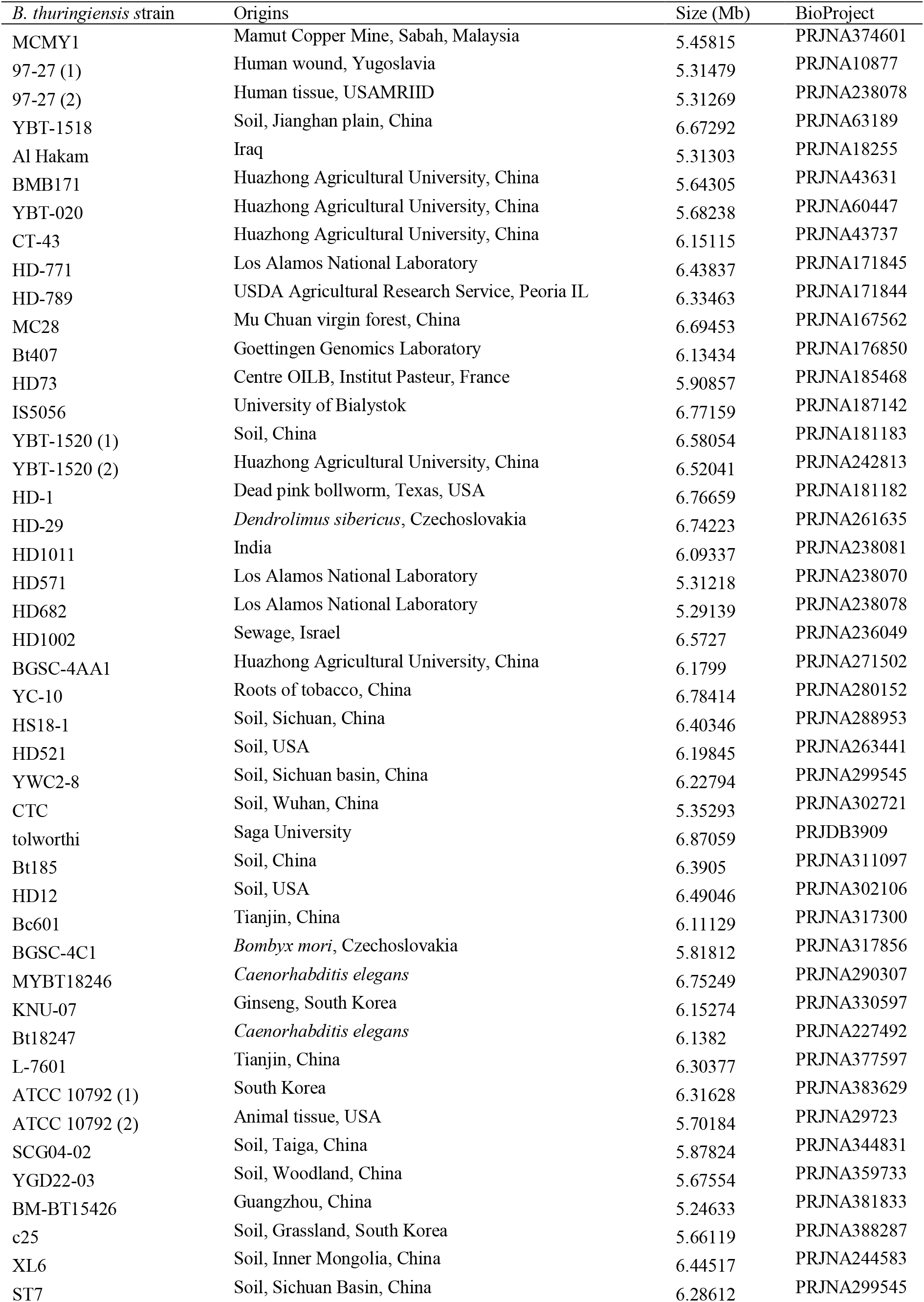
All *B. thuringiensis* genomes selected for comparison analysis.

## 3. Result and Discussion

### 3.1. Genome status of *B. thuringiensis*

There are a total of forty-four completed genomes of *B. thuringiensis* available in NCBI GenBank (as of, 2018). Three strains were found to be having two different complete genome submissions, which are the *B. thuringiensis serovar konkukian* strain 97-27, *B. thuringiensis* strain ATCC 10792 and *B. thuringiensis serovar kurstaki* strain YBT-1520. Both completed genomes of all three strains were included in the analysis for comparison purpose. All genomes of *B. thuringiensis* were inserted into the pipeline of QUAST version 4.6.3 for genome assembly quality assessment.

The Results was exported in HTML format and shown in Figure 1. Their genomes size varied between 5.20 (*B. thuringiensis* strain BM-BT15426) and 6.90 (*B. thuringiensis* serovar tolworthi) mega base pairs (MB) (Figure 1A). Figure 1B shows the largest contig length, L accounted for at least 80% to 100% of the bases of the assembly and Figure 1C shows the overall GC content of all genomes which was peaked at 35% approximately.

**Figure 1.**
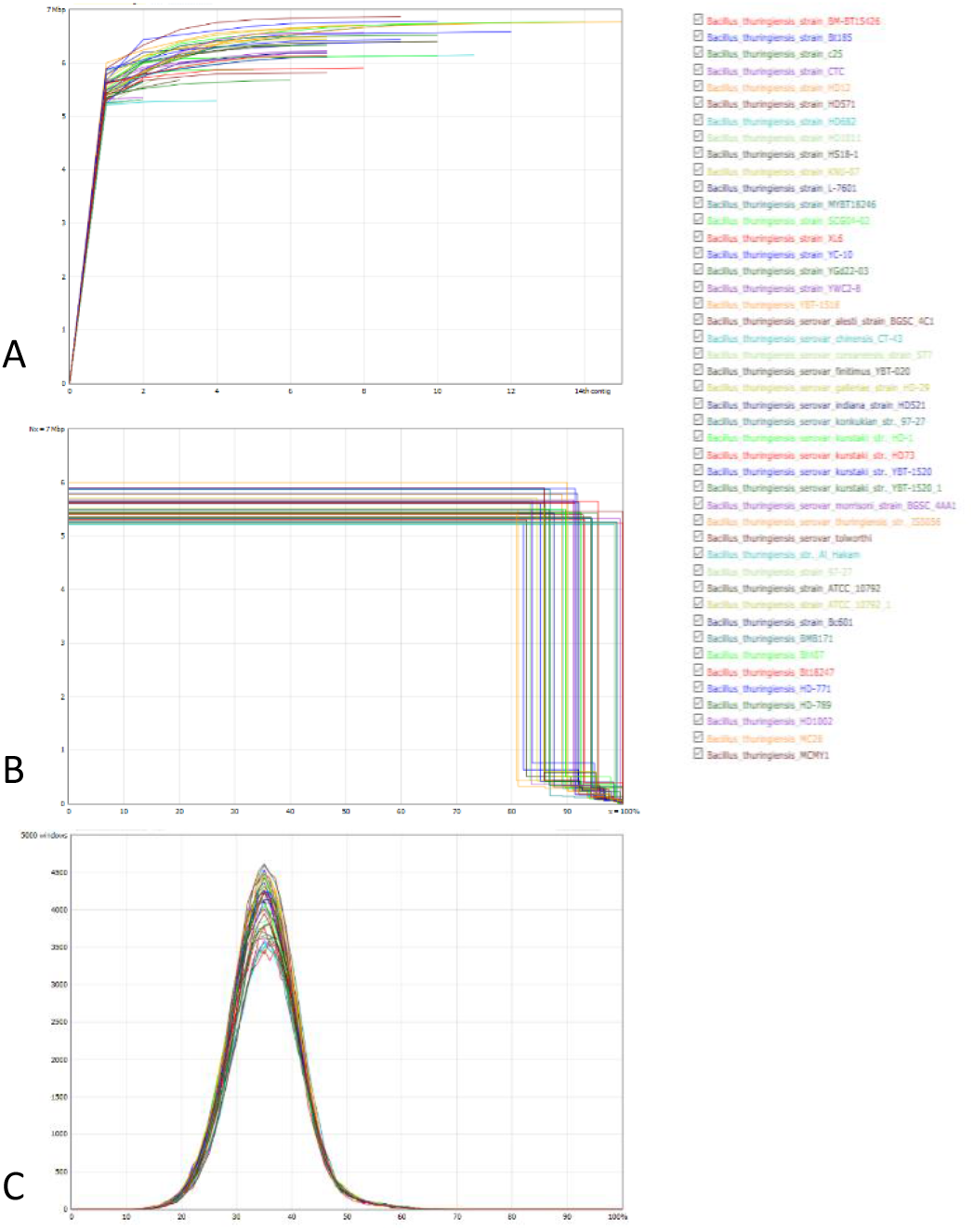
Genomes assembly assessment done by QUAST analysis.

### 3.2. Phylogenetic Information of *B. thuringiensis*

For phylogenetic reconstruction, *B. thuringiensis* genomes data were inserted into the Gegenees software version 2.2.1 together with a genome of *Escherichia coli* strain O157:H7 Sakai as a control for fragmented whole genome alignment and comparison analysis. *Escherichia coli* strain O157:H7 Sakai was presented with zero percent similarities with all *B. thuringiensis* strains. The target strain, *B. thuringiensis* MCMY1 matched the best with *B. thuringiensis* strain Bt407 and *B. thuringiensis* serovar thuringiensis strain IS5056 at 87%. Both genomes of the same strains, *B. thuringiensis serovar konkukian* strain 97-27, *B. thuringiensis* strain ATCC 10792 and *B. thuringiensis serovar kurstaki* strain YBT-1520 matched to each other in more than 99%. *B. thuringiensis* strain MC28 matched the least to all the other genomes at the lowest value of 45% with *B. thuringiensis* strain XL6. The fragmented whole genome alignment and comparison analysis was exported to SplitsTree software and converted into a phylogram based on the value of phylogenetic similarities. Figure 2 shows the phylogenetic tree generated with four mains clades were observed from the tree. Most of the *B. thuringiensis* strains were isolated from soil samples and insects body from all around the world, although some origins of the samples were not documented (Table 1). The target *B. thuringiensis* MCMY1 strain was classified in clade 3 and shared the same branch 3.3 with *B. thuringiensis* serovar thuringiensis strain IS5056, *B. thuringiensis* strain CT-43, *B. thuringiensis* strain Bt407 and both *B. thuringiensis* strain ATCC 10792 strains. Other members of Clade 3 are originated from China, USA and Germany. Clade 1 has the longest branches of all, this represents the group has the most diversified members among all.

**Figure 2.**
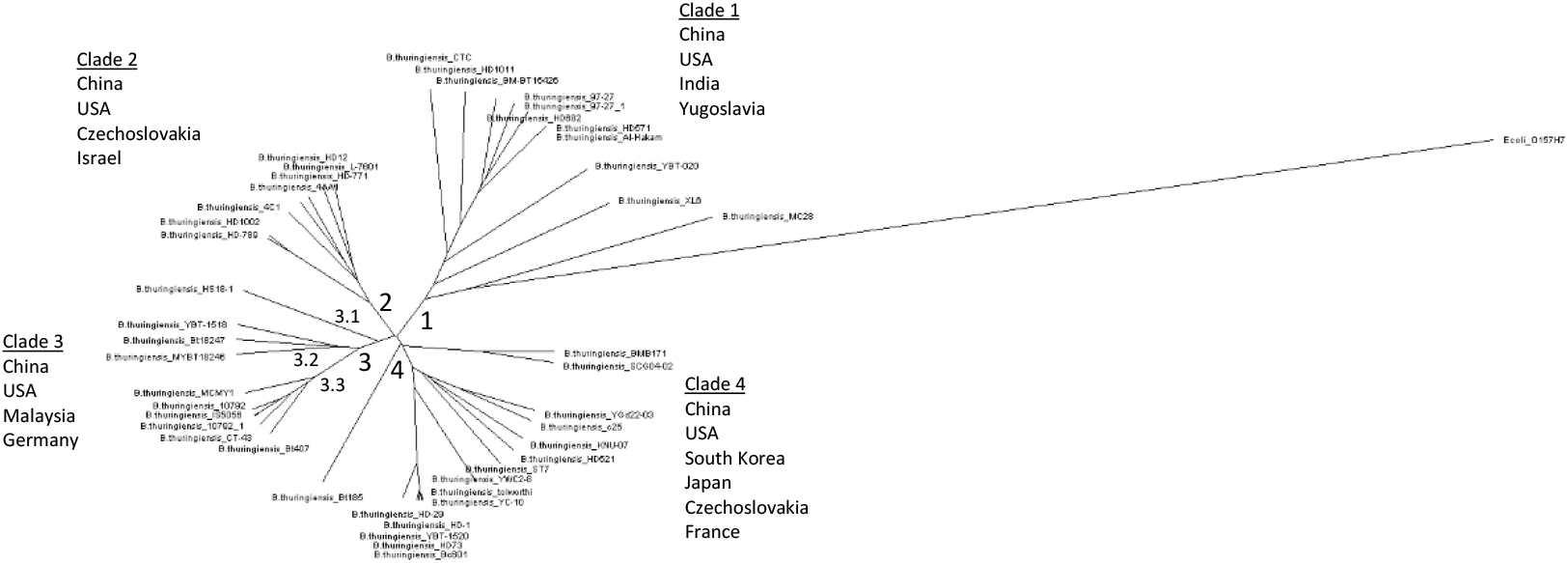
Phylogenetic tree generated for forty-five genomes of *B. thuringiensis* strains and their origin countries according to each clade.

### 3.3. Copper Homeostasis Genes of *B. thuringiensis*

The genomes data were imported to Rapid Annotations using Subsystems Technology (RAST) for annotations and the genes annotated to the subsystems of copper homeostasis, copper tolerance, copper uptake, copper transport system and blue copper proteins were retrieved from the database. Genes that are present in all genomes are the Copper-translocating P-type ATPase (CopA/B), Copper resistance protein C/D (CopC/D) and Conserved membrane protein in copper uptake (YcnI). YcnI is a protein that is speculated to be involving in copper acquisition and it is important during copper limiting conditions [15]. CopA/B is a type of protein that convey copper ions across cell surface and intracellular membrane [16], Cop C/D are the soluble periplasmic chaperone that bind to Cu(I) and Cu(II), they are hypothesized to be the transporter of essential copper through the inner membrane of cytoplasm [17]. The copper resistant CopABCD were reported from the *Pseudomonas syringae* pv. tomato genome [18]. Figure 3 shows the genes comparisons by using Blast Ring Image Generator (BRIG), all forty five genomes have shown high similarities in the CopABCD family together with the Copper-sensing transcriptional repressor (CsoR) and Copper (I) chaperone Z (CopZ). Only 23 genomes recorded the gene for Cytoplasmic copper homeostasis protein (CutC) and 19 genomes harbored the gene for Magnesium and Cobalt efflux protein (CorC). Figure 3 depicts the diversity of the the CutC and CorC genes in the forty-five *B. thuringiensis* genomes. The target genome *B. thuringiensis* MCMY1 harbors 11 genes except for Magnesium and cobalt efflux protein (CorC). CutC genes have been reported to bind to free copper radicals and reduce stress caused by copper contamination [19] while CorC gene plays a role in the transportation of Magnesium and Cobalt ions which has an indirect function to relieve the oxidative stress that caused by copper and other metal ions [20].

**Figure 3.**
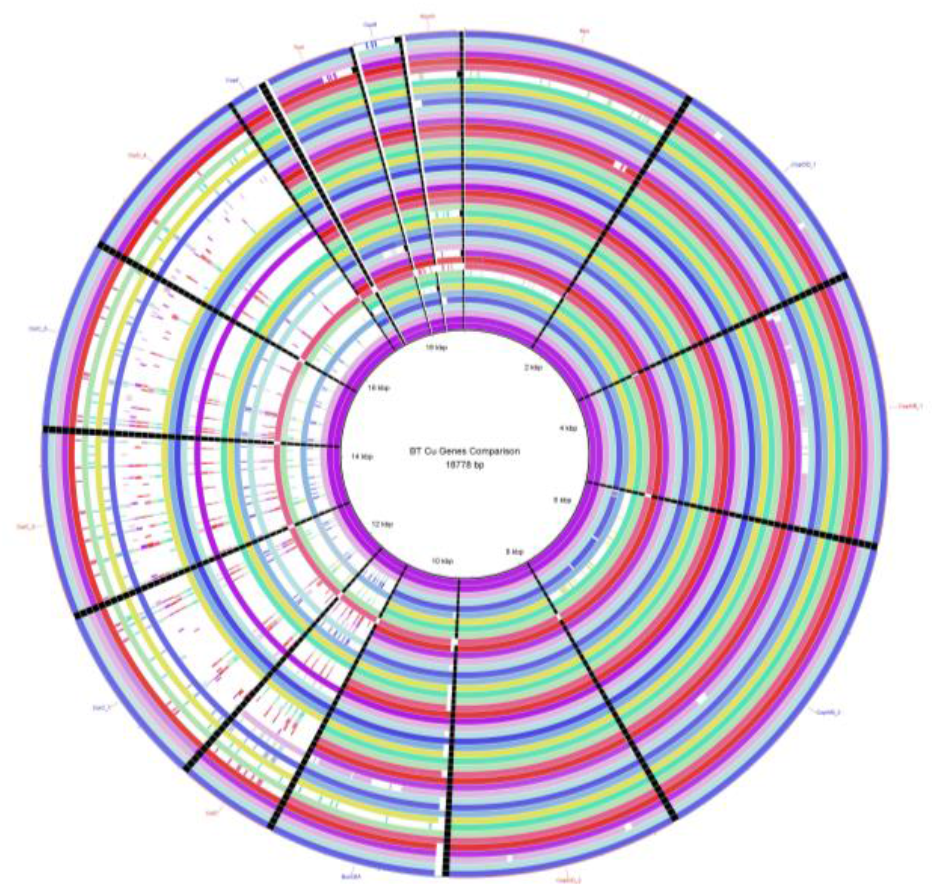
Copper genes comparisons by using Blast Ring Image Generator (BRIG).

As mentioned in the introductory section, some previous studies have stated that *B. thuringiensis* has been associated a wide range of environments and it can tolerate a different levels of copper induced stress [4,5,6,7,21,22]. However the degree of tolerance for copper recorded in their studies are contradistinctive. This can be caused by the unstandardized methodology practiced among the scientists or probably this can be an indicator to show that each *B. thuringiensis* strain that lives in a particular area have adapted to have its own ability to tolerate the copper contamination. Despite the lack of available publications related to the copper tolerance of *B. thuringiensis* in genetic aspect, the findings of current study suggest that Copper resistance genes family, CopABCDZ and its repressor (CsoR) may be the conserved genes in most of the *B. thuringiensis* genomes. However, the presence and the content of CutC and CorC genes in *B. thuringiensis* genome may play a role in altering the degree of tolerance to copper for each distinctive strain. Further investigations are required in order to support this hypothesis by understanding the expressions of these genes in *B. thuringiensis* during induced copper stress condition.

## 4. Acknowledgements

The authors are grateful for the financial support rendered by Ministry of Higher Education (Grant no. FRGS0455).

